# Neural Sampling Strategies for Visual Stimulus Reconstruction from Two-photon Imaging of Mouse Primary Visual Cortex

**DOI:** 10.1101/460659

**Authors:** Stef Garasto, Wilten Nicola, Anil A. Bharath, Simon R. Schultz

## Abstract

Deciphering the neural code involves interpreting the responses of sensory neurons from the perspective of a downstream population. Performing such a read-out is an important step towards understanding how the brain processes sensory information and has implications for Brain-Machine Interfaces. While previous work has focused on classification algorithms to identify a stimulus in a predefined set of categories, few studies have approached a full-stimulus reconstruction task, especially from calcium imaging recordings. Here, we attempt a pixel-by-pixel reconstruction of complex natural stimuli from two-photon calcium imaging of mouse primary visual cortex. We decoded the activity of 103 neurons from layer 2/3 using an optimal linear estimator and investigated which factors drive the reconstruction performance at the pixel level. We find the density of receptive fields to be the most influential feature. Finally, we use the receptive field data and simulations from a linear-nonlinear Poisson model to extrapolate decoding accuracy as a function of network size. We find that, on this dataset, reconstruction performance can increase by more than 50%, provided that the receptive fields are sampled more uniformly in the full visual field. These results provide practical experimental guidelines to boost the accuracy of full-stimulus reconstruction.

## INTRODUCTION

Neural firing patterns in primary sensory areas are commonly thought to contain information about the external world. For example, neurons in mouse primary visual cortex (V1) are thought to respond differently to natural and phase scrambled images [1]. To understand how such information is encoded in the neural responses, it is crucial to investigate how it can be extracted (decoded) from the firing patterns of populations of neurons[2]. While this endeavour has often taken the form of building a classifier to assign discrete categories to stimuli [1], the entirety of our sensory experience is not restricted to semantic categories, but extends to fine stimulus details. Previous work has attempted to address full-stimulus reconstruction by a variety of means. For example, Botella-Soler et al. [3] decoded artificial movies from the rat retina using nonlinear kernel regression, while Naselaris et al. used Bayes techniques to reconstruct natural movies from human functional Magnetic Resonance Imaging [4]. Linear decoders were used by Marre et al. [5] and Stanley et al. [6] to reconstruct low and high-dimensional stimuli from the salamader retina and the cat lateral geniculate nucleus, respectively.

Here, we investigate the problem of full reconstruction of natural stimuli from mouse V1 using two-photon calcium imaging recordings [7]. This problem poses several obstacles, including the low spatial acuity of mouse vision [8]. To the best of our knowledge, only one other paper has tackled the same challenge [9]. While Yoshida et al. [9] focus on how the information can be robustly represented by clusters of neurons, in this paper we analyze the influence of various factors, such as the neurons' Receptive Fields (RF) properties and the population size, on the reconstruction performance. We apply our decoder, an Optimal Linear Estimator (OLE) [10], to a public dataset and achieve an average frame-wise (pixel-wise) correlation coefficient of 0.28 (0.51), with a standard deviation of 0.26 (0.14). The frame-wise OLE accuracy is low, although still significantly better than chance. To improve performance, it is crucial to understand what features drive the reconstruction accuracy, and how should (future) data be collected to favour good decoding. Here, we show that RF density seems to be the most meaningful factor in explaining the quality of reconstruction. We also use the RF data to extrapolate to larger network sizes in simulations. Results suggest that, at least on this set of stimuli, decoding from more neurons could increase performance by 50% or more, depending on how the RFs are sampled. Due to the retinotopy of V1, our results imply that for full-stimulus reconstruction, experimental techniques should focus on sampling from larger areas of V1, rather than from more neurons in a patch.

## II MATERIALS AND METHODS

In this study, we used two main sources of data: a publicly available experimental dataset [11], and *in silico* simulations. Both involved the same set of stimuli and had matching RF and firing rates statistics. The *in silico* data was used to explore *what if* scenarios that could lead to higher performance.

### A. Data Collection

The data were recorded by Antolik et al. [11] and released under the terms of the Creative Commons Attribution Licence (https://creativecommons.org/licenses/by/4.0/). Briefly, the stimulus set is a collection of grey scale static images from David Attenboroughs BBC documentary *Life of Mammals.* Each image was presented for 500ms, interleaved by 1474ms of blank grey screen. Recordings were made at 7.6Hz and responses to an individual stimuli were computed as the average number of spikes across 5 consecutive two-photon imaging frames. A deconvolution algorithm [12] was used to infer spike counts for calcium traces. The final natural images used in the analysis are a patch of the displayed frames, centered around the location of the estimated population RFs, and down-sampled to 31 x 31 pixels. Details can be found in Antolik et al. [11]. Here, we used one of their imaging regions, containing 103 individual neurons. The training dataset, single-trial recordings, consists of 1800 image patches, while the test dataset, multi-trial recordings, has 50 image patches.

### B. Optimal Linear Estimator

We use a multi-input, multi-output linear estimator to decode the stimulus from the neural responses [10], [6], that is 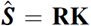, where 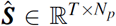 is the reconstructed stimulus, is the matrix of the linear filters and **R** ∊ ℝ^T × (N_n_ + 1)^ is the neural responses matrix. Here, *T* is the number of training frames, *N_p_* is the number of pixels in each frame and N_n_ is the total number of neurons. The extra column in the R matrix correspond to the bias term. Finally, the data in R is standardized. We estimate the optimal linear decoding filters by minimizing the reconstruction Mean Squared Error (MSE) with L_2_ regularization. The solution is given by the Optimal Linear Estimator (OLE) [10]: 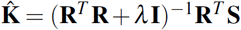. Here, ***S*** ∊ ℝ^T × N^_p_ is the matrix of (training) stimuli. The training dataset was used to estimate 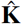, while performance is measured on the test set. A 5-fold cross validation was used to find the optimal value of X. Performance of the OLE was quantified using the correlation coefficient between target and reconstructed pixels (ρ_p_) or frames (ρ*f*). Neural response shuffling by randomly assigning the spatial pattern of neural responses between frames, was used to remove the input-output relationship between visual stimuli and neural activity. The chance level performance was measured with 400 repetitions of shuffling. To assess significance, a Wilcoxon signed rank test was used to test the hypothesis of equality of medians between shuffled and unshuffled conditions.

### C. Neural Response Models

We model the response r_ij_ of an individual neuron *j* to a single stimulus frame *i* is given by r_ij_ *j* = **L(s^i^)h**^j^. In the linear model, L**(S**^i^) = s*i* and 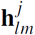 is obtained using a Spike Triggered Average with Laplacian regularisation [11]. That is, 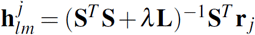, where **L** is a discrete Laplacian operator and r_j_ ∊ ℝ^T × 1^ the full response of neuron *j.* In the Pyramid Wavelet Model (PWM) [13], [4], L **(S** ^i^) ∊ ℝ^1 × N_F_^ is a nonlinear transform that consists in first projecting the stimulus frame onto a set of Gabor filters with different frequencies, locations, phases and orientations, followed by a point nonlinearity (either a ramp function or the sum of square from quadrature-phase wavelets, to model both simple and complex cells). *NF* is the number of Gabor wavelets considered (here, 33990). Each entry 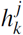 of the weight vector 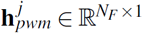 quantifies how much the response of neuron *j* depends on feature *k*. Such a vector was estimated using the L_2_ boost algorithm with early stopping (via the open source STRFLab toolbox [14]), to encourage sparseness [13].

### D. Receptive Field Estimate

For neurons that were well predicted by the PWM (correlation coefficient between measured and predicted responses higher than 0.3), a linearized version of the neuron’s RF was obtained by the sum of all the wavelets used in the PWM, weighted by the vector 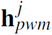. Otherwise, the RF computed with the linear model 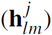 was used, if the model predicted the neuron’s response with a correlation coefficient better than 0.3. The neurons that fell below both thresholds did not contribute a RF to the following analysis. Then, a single elliptical Gabor function was fit to each RF with a Gaussian envelope taken as the boundary. The orientation of the envelope (θ) was considered to be the preferred orientation of that RF. If the fit failed, and the neuron exceeded the PWM threshold, the Gaussian envelope of the base wavelet corresponding to the highest entry in the weights vector 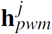 of the PWM was used. Otherwise, that RF was dropped. A pixel in a 31 × 31 frame was considered to belong to a RF if it was within that RF’s boundary. The RF coverage of a pixel is the number of RFs that included that pixel.

### E. Regression Model of Pixel-wise Performance

We built a multivariate linear regression model to predict the pixel-wise performance of the OLE (p_p_) from 7 different variables. Three of these features describe the ensemble of RFs covering each pixel, specifically its cardinality and its heterogeneity. The latter is given by the average and the spread (for circular variables) of the preferred orientations of all the RFs in the ensemble. The other 4 features are the mean, standard deviation, skewness and kurtosis of the distribution of intensity values at a particular pixel location across the test dataset. Dependent and independent variables were standardized before fitting. We used permutation importance to quantify the relative contribution of each regressor to the OLE performance. For each feature, we shuffled its values and then evaluated the relative change in the adjusted r-squared of the regression model. We report the mean relative change in performance for each regressor across 1000 repeats, normalized to a total of 1. Error bars are the standard deviations over the repetitions, normalized by the same amount. We also used two different sparsity inducing algorithms-stepwise linear regression (according to Bayesian Information Criterion) and the LASSO (regularized elastic-net with alpha = 0.95)-to check which regressors would survive feature selection. We used the 1-Standard-Error rule to select the final model.

### F. In silico Neurons Generation

To generate *in silico* data, we first extracted the RFs parameters (locations, size, orientations, spatial frequency, phase, amplitude and bias) from the Gabor fits described in the previous section. Then, we created new RFs by sampling from these distributions. We simulated two different experimental conditions. In the “worst case” scenario, locations for the simulated RFs were drawn from the experimental distributions. In the “best case” scenario, we augmented the experimental distributions of RF locations with shifted replicas of itself. Each shift was randomly drawn and correspond to shifting the RFs in the visual field. The two cases coincide when we simulate 100 neurons. The sampling of all the other parameters was the same in both scenarios. Neural responses were generated using a linear-nonlinear Poisson model: 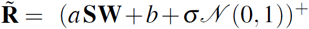. Here, 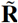 are the *in silico* responses, W are the simulated RFs, σ is a scalar (set to 2.7 to match OLE accuracy when using 100 synthetic neurons), *N*(0,1) is the standard normal distribution, and (.)+ is the ramp function. Furthermore, *a* and *b* are parameters whose distributions we estimated from the data using a least-squares fit on the experimental responses and then sampled during the simulations. Neurons that exceeded a maximum firing rate (set by experimental data) were discarded. Responses were generated to the same set of stimuli used experimentally and with the same protocol of single-and multi-trial responses for training and testing data, respectively.

## III. RESULTS

### A. A Simple Linear Secoder Can Reconstruct Natural Images from Mouse Primary Visual Cortex

Despite its simplicity, a linear decoder has been shown to achieve good performance [10], [6], [5] and is compatible with highly nonlinear encoding mechanisms [2]. Here, we used an OLE to reconstruct full visual stimuli from two-photon imaging of mouse V1 (layer 2/3). The optimal linear decoding filters were computed to minimize the MSE of the reconstruction, subject to L_2_-regularization. The relative weight of the regularization term was computed using crossvalidation. Performance was quantified through the Pearson correlation coefficient both between each target and reconstructed pixel (ρ_p_) and between each target and reconstructed frame (ρ*f*). The full distribution of p*f* is shown in Fig 1a, while that of ρ_p_ is shown in the upper histogram of Fig 3a. The mean values (standard deviations) are 0.28 (0.26) and 0.51 (0.14) for ρ*f* and ρ_p_, respectively. Some, but not all, of the frames are predicted with good accuracy (see Fig 1b), although the distributions are spread across a broad range of values. Similarly, some pixels are better decoded than others (Fig 1c). For completeness, the average MSE between targets and predictions across frames and pixels is 0.07 (with a standard deviation of 0.07). To evaluate whether the OLE performed better than chance, we used a neural shuffling procedure to destroy the input-output relationship between the responses and the stimuli and tested the null hypothesis of chance level performance using a Wilcoxon-signed rank test. The chance level distribution of ρ*f* is shown in Fig 1a. The *p*-values obtained comparing both correlation coefficient distributions to their chance level counterparts are lower than 10^-8^: the OLE performance are significantly better than chance.

**Fig.1.**
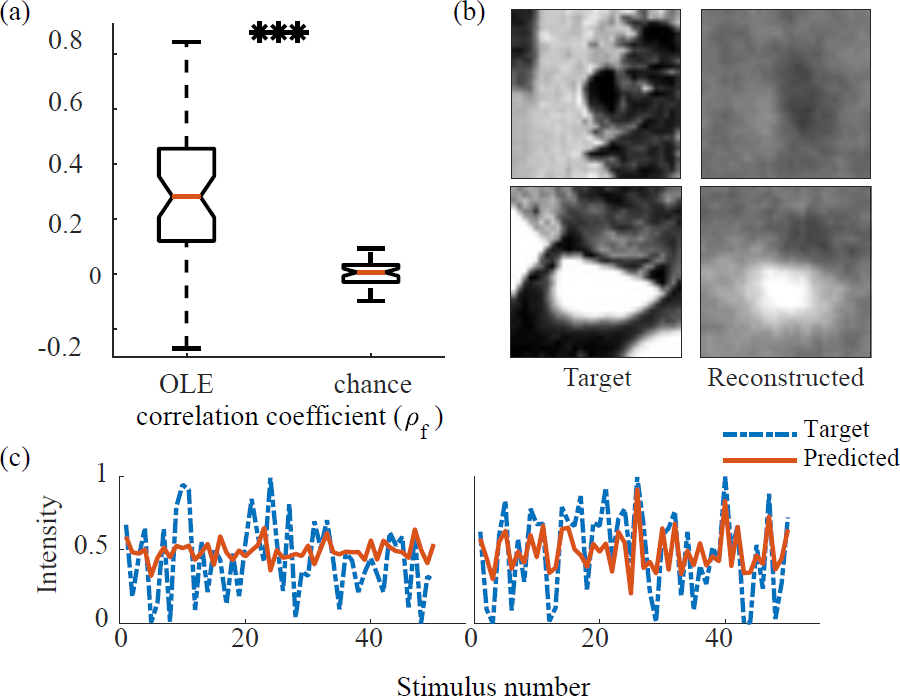
Linear decoder performance. (a) Performance of the linear decoder across frames, compared with chance level performance. The presence of three asterisks imply that the OLE is significantly better than chance with p < 0:0001 (Wilcoxon signed rank test). (b) Two examples of reconstructed stimuli, at different performance levels. The colormap range for all frames is between 0 (black) and 1 (white). (c) Two examples of reconstructed pixels (red solid lines) with their respective targets (blue dashed lines). Left and right pixels are taken from the stimulus upper left corner and center, respectively.

### B. Receptive Field Coverage Explains Reconstruction Accuracy

We computed the RFs for the whole population by fitting a single Gabor filter to the linearized RF obtained after fitting each neuron with a pyramid wavelet model. If the model did not predict the neural response sufficiently well, a laplacian regularised linear model was used, instead. Fig 2a shows all the RFs from the population superimposed on the stimulus visual field. Furthermore, Fig 2b shows the level of RF coverage for each individual pixel, normalized between 0 (black) and 1 (white). The higher the coverage, the more neurons had a RF localized around that spatial location. It can be seen that the RFs cluster around the center of the image, consistently with the experimental protocol described by Antolik et al. [11]. Two examples RFs are displayed in Fig 2c (right column). In the left column, instead, we show the corresponding decoding filters for the same two neurons. It can be seen that the linear filters of the encoding and the decoding model have a considerable overlap, even though they have been optimized independently. However, not all neurons showed a similar result, especially those with more noisy decoding filters.

**Fig.2.**
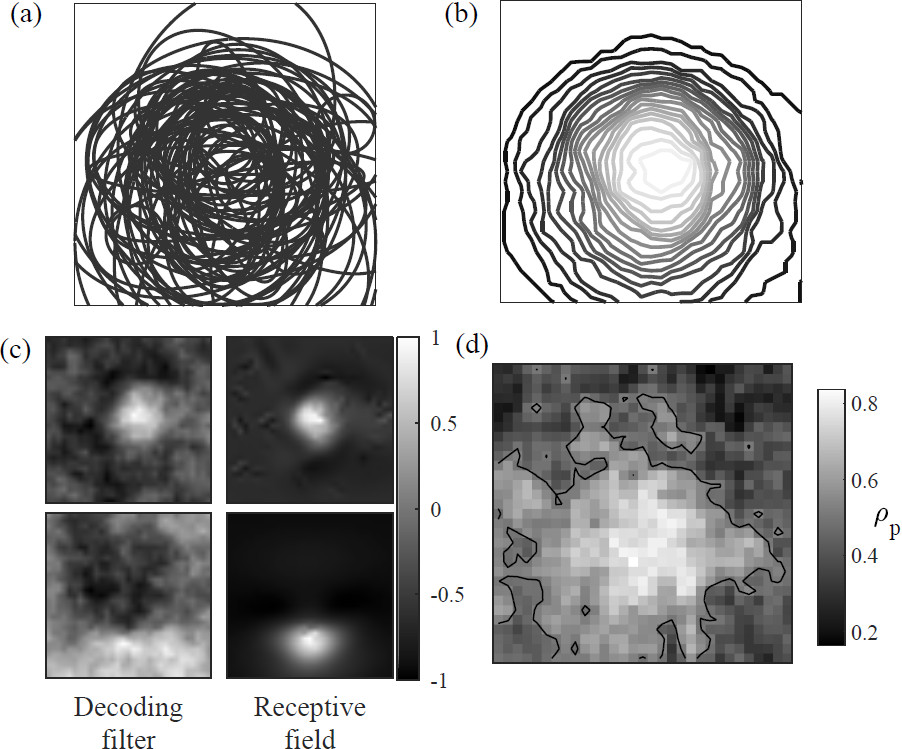
Neurons' receptive fields and OLE performance. (a) All estimated receptive fields from the population. (b) Receptive fields coverage (normalized between 0, black, and 1, white) as a function of the spatial location. (c) Two examples of receptive fields and decoding fields from the same neuron. (d) Linear decoder performance (ρp) as a function of spatial location.

The clustering of the RFs in the middle of the stimulus is likely to increase the information content around the central pixels carried by the whole neural population. Indeed, we found a strong positive relationship between the performance of the algorithm at each pixel, and the RF coverage, similarly to that reported by Botella-Soler et al. [3] (Fig 2d and 3a, correlation coefficient of 0.79). However, while the RF coverage is likely to explain a large proportion of the OLE accuracy, a significant role could also be played by other factors, such as the statistics of each individual pixel which, given the small sample size of the test dataset, are likely to be different. To test for this, we computed 6 other potentially relevant pixel-wise features: the mean and the spread of the orientation of the local RFs, and the mean, standard deviation, skewness and kurtosis of the pixel intensity levels. We then built a multivariate linear regression model (MLR) to predict ρ_p_ from the RF coverage and the 6 quantities above. Being only interested in the fluctuations around the mean, we standardized both the independent and the dependent variables, leading to a null intercept. The results from the model (coefficients are presented with their confidence intervals) are shown in Table I (first column, the adjusted R-squared is 0.67), where asterisks indicate statistically significant regressors. We then used the permutation importance technique to quantify the relative contribution of each feature to the model performance: the outcome is reported in Fig. 3b (bar heights are normalized so that their sum is 1). Both Table I and Fig. 3b show that the RF coverage is the feature with the most influence on the reconstruction performance, followed by the standard deviation of the pixel intensities and the spread of the RF orientations. Finally, we verified that the results were insensitive to the regression technique by using stepwise and LASSO linear regression (see Table I). These two procedures return the minimal set of features needed to explain the reconstruction performance.

**Table 1.** MULTIVARIATE LINEAR REGRESSION OF THE OLE PERFORMANCE AGAINST RFS AND IMAGE FEATURES.

|  | MLR | step MLR | Lasso |
| --- | --- | --- | --- |
| RF coverage | $0.75 \pm 0.04^*$ | $0.74 \pm 0.04^*$ | $0.71 \pm 0.03^*$ |
| mean ($\theta$) | $-0.02 \pm 0.04$ | 0 | 0 |
| spread ($\theta$) | $0.11 \pm 0.04^*$ | $0.11 \pm 0.04^*$ | $0.08 \pm 0.04^*$ |
| mean | $-0.04 \pm 0.07$ | $0.06 \pm 0.04^*$ | $0.03 \pm 0.04$ |
| st. deviation | $0.17 \pm 0.06^*$ | $0.17 \pm 0.04^*$ | $0.14 \pm 0.04^*$ |
| skewness | $0.14 \pm 0.07^*$ | 0 | 0 |
| kurtosis | $0.13 \pm 0.06^*$ | 0 | 0 |
| adj. R-squared | 0.67 | 0.67 | 0.66 |

**Fig.3.**
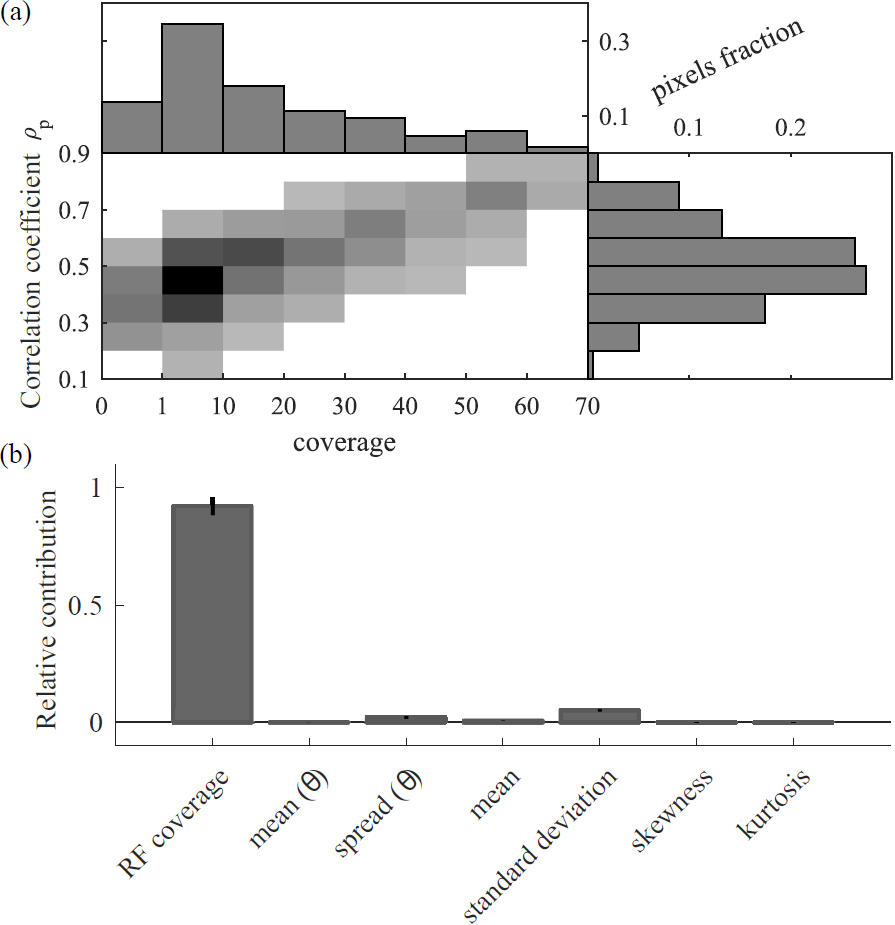
Explaining the OLE performance. (a) Accuracy of the linear decoder (ρ_p_) as a function of the receptive field coverage, and respective histograms. (b) Relative contribution of each feature to the performance of the linear decoder (ρ_p_), using a linear regression model.

### C. Effect of Increasing the Number of Neurons

Recording from a larger number of neurons would consequently increase the RF coverage and, thus, reconstruction performance. We investigated how much of an improvement could be obtained by generating populations of *in silico* neurons of different sizes, yet with RF and firing statistics extrapolated from the existing data. We tested two different scenarios: one that corresponds to increasing the neural density of the imaged cortical area (“worst case”), and one that is akin to recording from more cortical areas, all with fixed neural density (“best case”). Both cases result in more neurons being decoded. Specifically, we simulated populations between 100 and 4000 neurons, with RF characteristics similar to those of the recorded cells as confirmed by computing the RF coverage (normalized between 0 and 1) for 100 *in silico* neurons (Fig. 4a, on the left, to be compared with Fig. 2b). Gaussian noise was added to match the experimental performance. However we could only match ρ*f*, while ρ_p_ was always higher (0.66 on average). The mean firing rates of the simulated neurons were also generally higher than for the experimental ones, likely a consequence of the added noise.

Results for different population sizes are reported in Fig. 4b as the mean and the standard deviation across 20 simulations of the average ρ *f*. In both scenarios performance improve considerably, quantitatively and qualitatively (example reconstructions are also shown in Fig. 4b for various numbers of neurons, color-coded according to the legend in the plot on top): increasing the population size means that more features of the stimuli are captured by the reconstructions. Furthermore, for the “worst case” scenario, accuracy seems to saturate around 2000 neurons at an average of ρ*f =* 0.48 (71% increase), while the “best case” line shows an always positive slope, with p*f =* 0.61 (an increase of more than 100%) for 4000 neurons and pf = 0.49 already reached with 500 neurons. The reason for the improved reconstruction accuracy is likely due to a more uniform coverage of the visual field by the simulated RFs (Fig. 4a, on the right).

**Fig.4.**
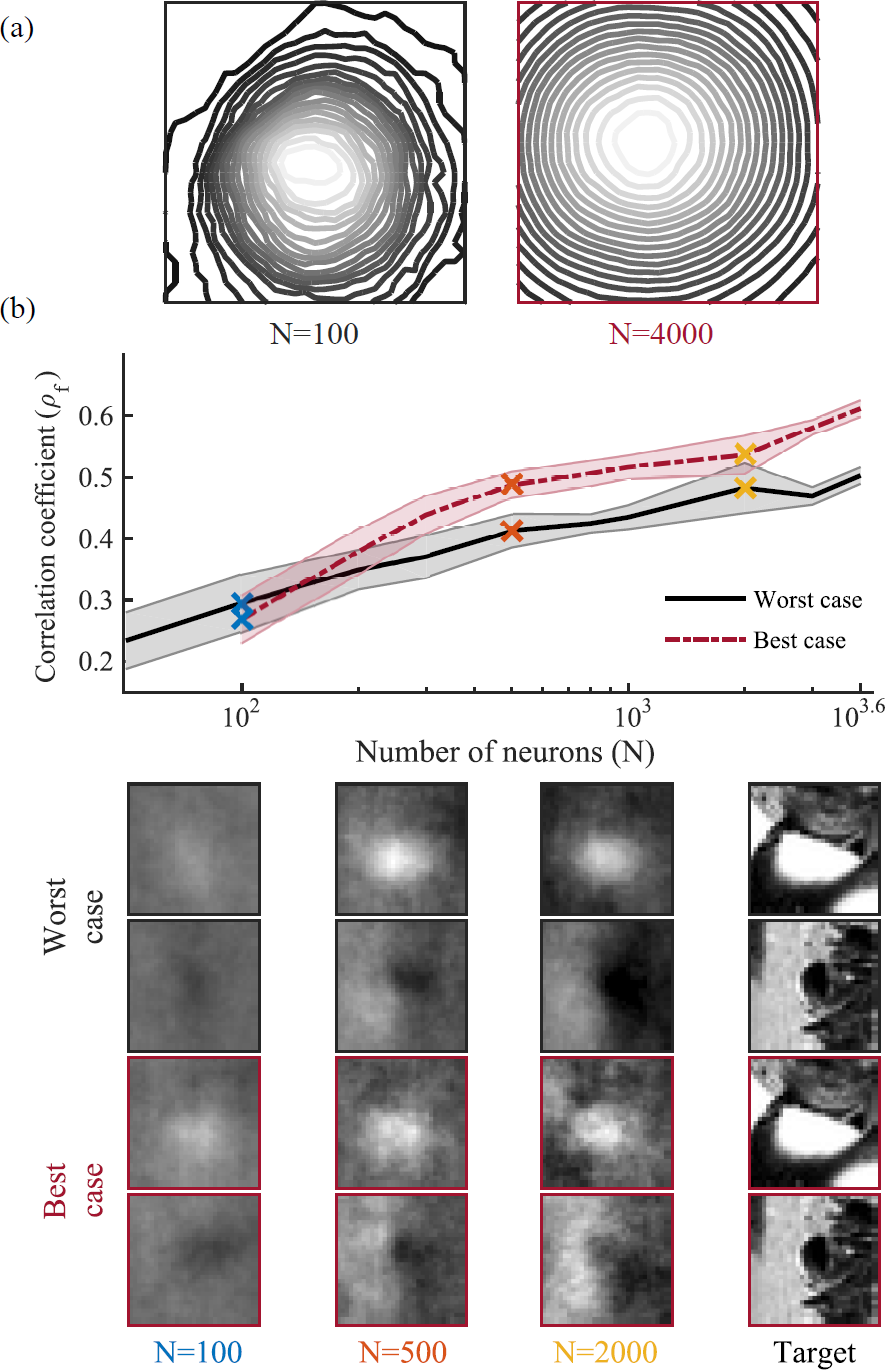
Effect of increasing numbers of neurons. (a) Receptive fields coverage (normalized between 0, black, and 1, white) as a function of spatial location for 100 (left) and 4000 *in silico* neurons (right, under the “best case” scenario). (b) Top: semi-log plot of reconstruction performance versus population size for “best” (red dashed line) and the “worst case” (black solid line) scenarios. Lines and shaded area are the average and the standard deviation of the accuracy across 20 simulations. Bottom: example reconstructions for various numbers of neurons, color-coded according to the legend in the semi-log plot.

## IV. DISCUSSION

In this paper, we used a linear decoder to reconstruct full visual stimuli from two-photon imaging recordings in layer 2/3 of mouse V1. The results obtained are significantly better than chance, although there can be large difference in accuracy across frames and pixels. We showed that the local RF coverage was the best predictor for the pixel-wise OLE performance (as hinted by the strong correlation between the two features [3]), followed by the standard deviation of the pixel intensities and the heterogeneity of the RFs. Using simulations from a linear-nonlinear Poisson model, we computed the increase in performance accuracy that could be obtained with larger populations sizes, in two possible experimental scenarios. Confirming the regression results, we report a higher accuracy for the case where RFs are more uniformly distributed across the visual field. While we expect the actual increase in performance to fall somewhere between the “worst” and the “best case” scenario, the reported results could be used to guide future experimental procedure aimed at full visual stimulus reconstruction. For future work, we aim at improving the simulations with a more realistic noise model. Furthermore, it remains to be seen whether a nonlinear decoder [15] could improve reconstruction performance for the experimental or the simulated data.

